# Programmable DNA Origami Caps for Site-Selective Functionalization of Microtubule Tips and Lattice Defects

**DOI:** 10.64898/2026.05.08.722927

**Authors:** Henry Carey-Morgan, Brenda Palestina-Romero, Azra Atabay, Jonathan Bath, Andrew Turberfield, Elisha Krieg, Stefan Diez

## Abstract

Microtubules are central components of cytoskeletal transport systems and have been widely repurposed as active elements in motor-driven nanodevices. However, site-specific functionalization of stabilized microtubules remains a fundamental challenge, as the tubulin lattice presents chemically indistinguishable binding sites along its length. Here we report a strategy for selective end-functionalization of stabilized microtubules using DNA origami nanostructures. By coupling DNA origami to Fab fragments targeting acetylated α-tubulin Lys40 within the microtubule lumen, and exploiting steric exclusion of the origami from the lattice interior, binding is confined to accessible sites at microtubule ends and lattice defects. Using a six-helix bundle origami as a minimal construct, we demonstrate selective tip labelling of gliding microtubules without perturbing kinesin-driven motility. The same structures additionally mark lattice defects, enabling dynamic visualization of defect sites during transport. Furthermore, we show that tip-bound origami can hybridize with complementary DNA strands to capture cargo from surfaces in motion, establishing programmable, end-specific loading. This approach introduces a generalizable route to spatially controlled functionalization of cytoskeletal filaments, enabling new capabilities in molecular transport, nanoscale assembly, and the study of microtubule integrity and repair.

## Introduction

Microtubule-kinesin gliding assays have emerged as powerful platforms for nanoscale transport, sensing, and computation, enabling the use of cytoskeletal filaments as active components in engineered nanodevices. In particular, nanodevices engineered around gliding assays have been used for example to map nanoscale surface-topographies [1], detect analytes in solution [2-8], discover, collect and sort cargo from a surface [9-20], or decode mathematical problems in biocomputational devices [21-23]. Despite this versatility, a central limitation persists: the lack of spatial control over microtubule functionalization. Both pre-polymerization modification of tubulin and post-polymerization conjugation strategies [24] produce uniformly decorated filaments, as stabilized microtubules present chemically indistinguishable tubulin subunits along their length. This absence of site specificity restricts the development of more sophisticated, addressable transport systems.

In living systems, the microtubule lumen serves as a distinct biochemical compartment targeted by proteins that regulate filament stability and post-translational modification [25-28]. One such modification, acetylation of α-tubulin at lysine-40 (αK40), is located on a luminal loop accessible to binding proteins [29-31]. Antibodies raised against this modification can recognize tubulin within the microtubule lumen [32-35]; however, because acetylated residues are distributed throughout stabilized microtubules, such recognition alone does not confer spatial selectivity [36,37].

Here we address this challenge by introducing a steric gating strategy that restricts luminal binding to microtubule ends. We couple Fab fragments targeting acetylated α-tubulin to DNA origami nanostructures whose dimensions exceed that of the microtubule lumen. This design prevents the antibody-origami complex from penetrating deeply into the lattice, effectively confining binding to accessible sites at filament ends or lattice discontinuities. In this way, DNA origami acts as a programmable nanoscale cap that converts a non-specific luminal epitope into a site-selective binding mechanism.

DNA origami provides a uniquely versatile platform for nanoscale engineering, enabling precise control over geometry, valency, and chemical functionality [38-45]. By integrating these capabilities with microtubule-based transport, the resulting structures can be tailored to perform specific tasks at defined filament locations. This approach establishes microtubule ends as addressable functional sites, analogous to sockets that can be equipped with interchangeable nanoscale modules.

## Results

To achieve site-selective functionalization of stabilized microtubules, we designed a DNA origami-antibody construct that exploits steric exclusion within the microtubule lumen. A six-helix bundle DNA origami was chosen as a simple, rigid scaffold. Although not geometrically matched to the lumen as a true “plug,” its size and negative charge were expected to hinder deep penetration into the ∼15 nm inner diameter of the microtubule, thereby restricting binding to accessible regions.

The origami structure was engineered with 22 single-stranded overhangs for hybridization of Atto647N-labeled oligonucleotides, enabling fluorescence visualization, and three additional overhangs for attachment of antibody fragments (**Fig. 1A**). Folding was achieved using a standard M13 scaffold and staple strands, and successful assembly was confirmed by agarose gel electrophoresis and spectrophotometric analysis (**Fig. S1**).

**Figure 1:**
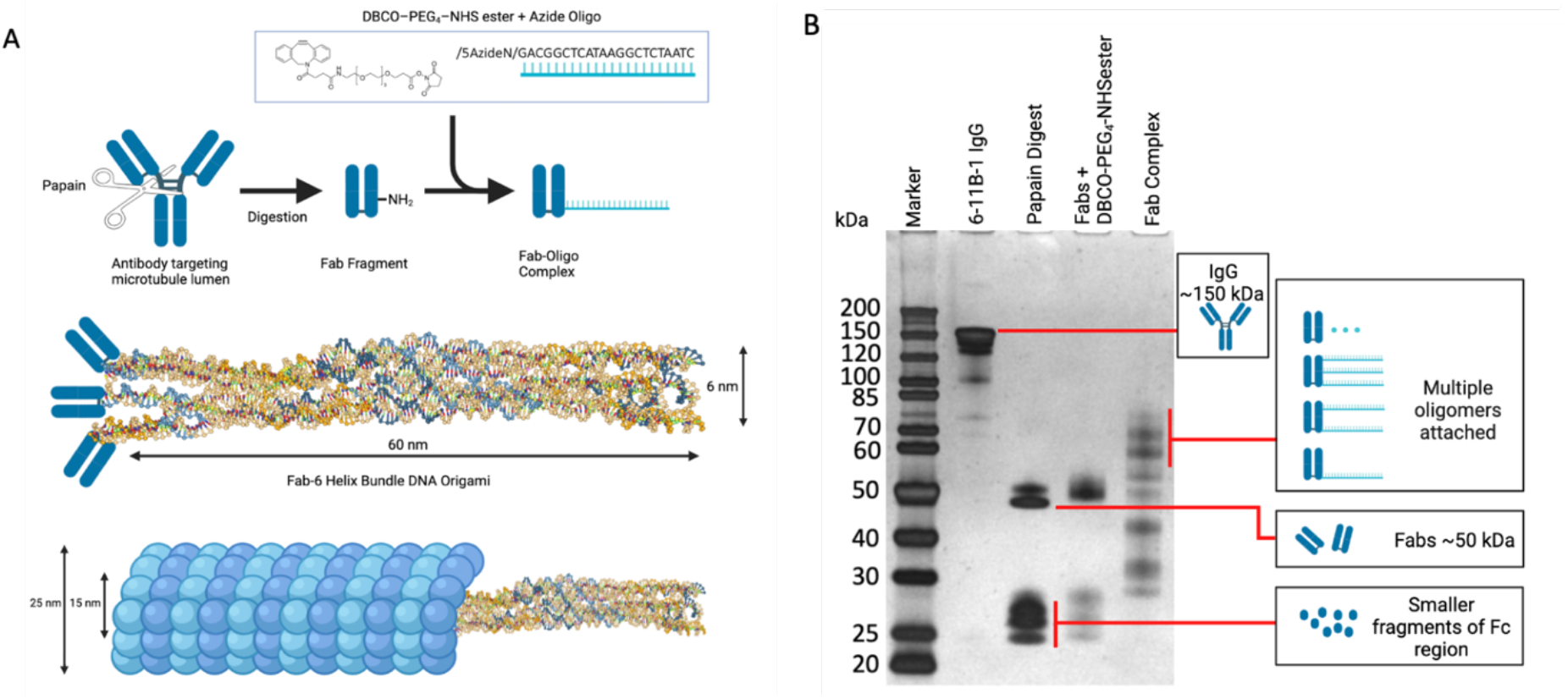
Design and assembly of DNA origami–Fab complexes for sterically gated luminal binding. **A)** Illustration showing the bioconjugation scheme. **B)** Silver-stained SDS-PAGE of 6-11B-1 antibody fragmentation and Fab Complex formation. The full antibody (6-11B-1 IgG, MW ∼150 kDa) is cleaved by the papain digestion into Fab and Fc fragments which are further digested (Papain Digest). The addition of the DBCO-PEG_4_-NHS ester crosslinker (400 Da) causes a slightly raised and smudged band as multiple crosslinkers may bind to a single Fab (Fabs + DBCO-PEG_4_-NHSester). Addition of azide functionalized DNA oligomers (∼6 kDa) then causes the separation of the Fab + crosslinker into multiple bands clearly delineating the binding of multiple oligomers (Fab Complex).

To target the microtubule lumen, monoclonal anti-acetylated α-tubulin antibodies (clone 6-11B-1) were enzymatically digested into Fab fragments, reducing steric bulk and enabling multiple fragments to access the lumen simultaneously [46]. The Fab fragments were conjugated to azide-functionalized DNA oligonucleotides via DBCO-mediated click chemistry [47]. SDS–PAGE analysis revealed successful conjugation, with discrete band shifts corresponding to Fabs bearing one or more DNA strands (**Fig. 1B**; **Fig. S2**). These Fab-DNA complexes were then hybridized to the origami scaffold, forming multivalent DNA origami-Fab assemblies, which were further verified by transmission electron microscopy (TEM) (**Fig. S3**). Following incubation with taxol/GMPCPP-stabilized microtubules, these constructs were expected to bind luminal acetylation sites while being sterically restricted from diffusing deep into the lattice, thereby forming effective “caps” at accessible sites.

To test whether steric exclusion results in preferential end binding, DNA origami–Fab complexes were incubated with stabilized microtubules and introduced into kinesin-driven gliding assays. Fluorescence imaging revealed clear localization of Atto647N-labeled origami at microtubule tips (**Fig. 2A,B; Supplementary Movie 1**), demonstrating successful capping. Capping yields of approximately 5% were obtained under the conditions used. In control experiments, DNA origami lacking Fab conjugation showed no detectable binding across hundreds of observed microtubules, confirming that attachment is mediated specifically through antibody recognition of luminal acetylation sites.

**Figure 2:**
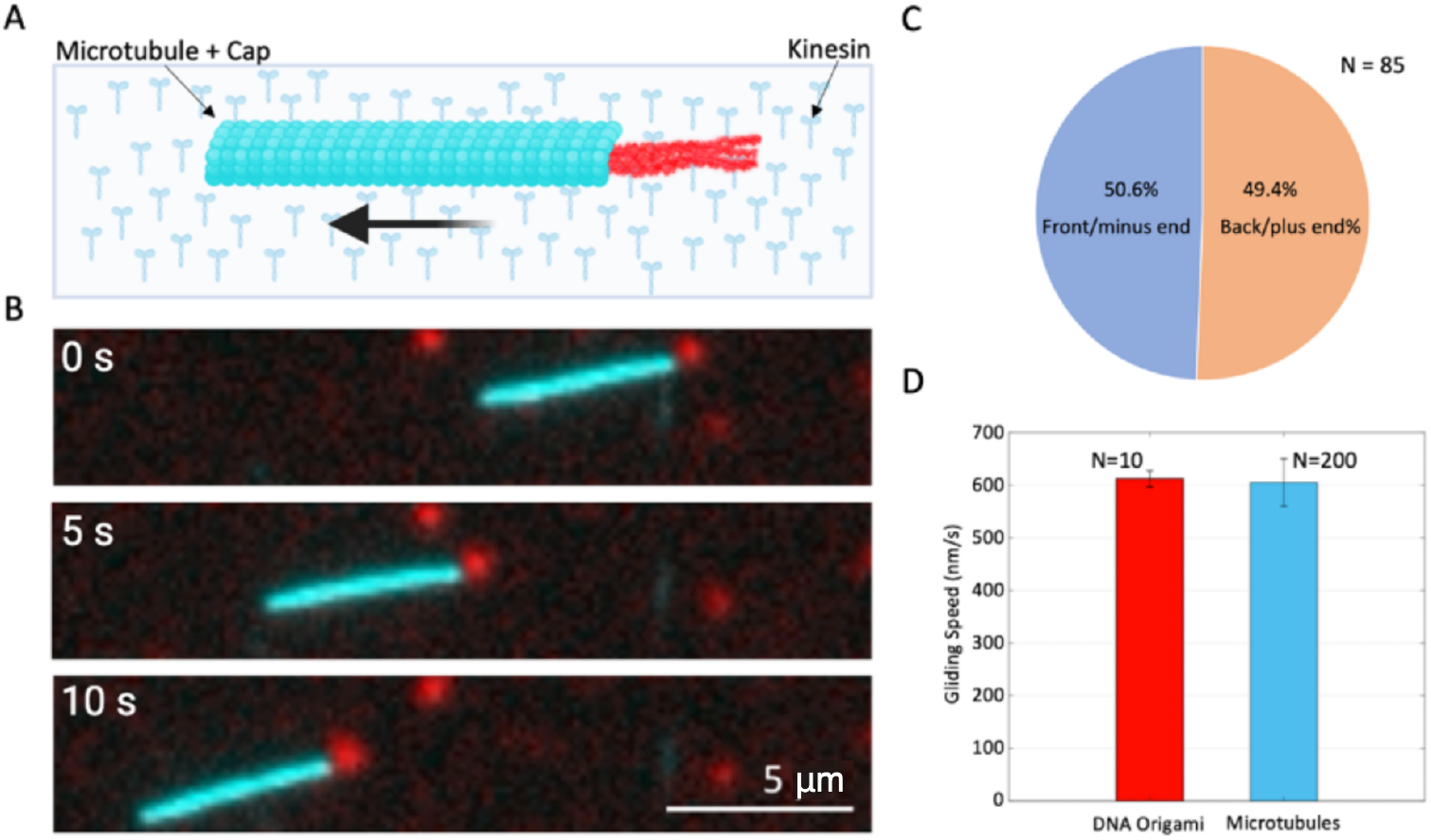
Selective microtubule end capping and preservation of motility. **A)** Schematic of six-helix bundle labelled with Atto647N (red) capping a TAMRA labelled microtubule (cyan) gliding on surface-immobilized kinesin motors. **B)** Fluorescence microscopy images of TAMRA labelled microtubule (cyan) with Atto647N DNA Origami cap. The Origami appears to trail after the microtubule as imaging in each fluorescent channel is sequential, not simultaneous. Unbound DNA Origami can be seen adhered non-specifically to the surface. **C)** 85 microtubules were observed with tip-adhered DNA Origami caps almost exactly even split between caps bound to the ‘front’ or ‘back’ of the microtubules. **D)** The speed of ten microtubules with DNA Origami caps was measured utilizing the tracking software FIESTA and there was no difference in their gliding speed compared with ∼200 unmarked microtubules. A Welch’s two-sample t-test indicated no significant difference in gliding speed between DNA origami and microtubules (p ≈ 0.78).

Several key observations were made regarding the behavior of capped microtubules. First, caps exhibited no directional preference, appearing at both leading and trailing ends with approximately equal frequency (**Fig. 2C**). This indicates that acetylated αK40 residues are accessible at both microtubule termini and that the capping mechanism does not distinguish between plus and minus ends. Second, the presence of the cap did not measurably affect kinesin-driven motility. Quantitative analysis of gliding velocities showed no significant difference between capped and uncapped microtubules (**Fig. 2D**), indicating that the origami structure does not act as a steric obstacle to motor progression along the lattice. Together, these results demonstrate that DNA origami-Fab constructs can selectively functionalize microtubule ends while preserving their transport properties.

In addition to tip localization, a fraction (∼11%) of DNA origami structures (n = 172) was observed bound along the microtubule lattice (**Fig. 3A**). We hypothesized that these events arise from access to the lumen through structural defects in the microtubule lattice (**Fig. 3B**). To test this, the motion of lattice-bound origami was analyzed during gliding. This structure exhibited a periodic lateral displacement consistent with the helical rotation of microtubules, with a measured pitch of ∼8 μm (**Fig. 3C**) [48]. This behavior indicates that the origami remains externally attached and rotates with the filament, rather than diffusing freely within the lumen. TEM provided direct structural evidence, revealing DNA origami associated with discontinuities in the microtubule lattice (**Fig. 3D**). Furthermore, microtubules aged over several months exhibited a markedly higher frequency of lattice-bound origami, consistent with an increased density of defects arising from lattice degradation over time (**Supplementary Movie 2**). These observations support the interpretation that DNA origami–Fab complexes can access the lumen at defect sites and subsequently bind to acetylated tubulin. As such, the caps provide a means of directly labeling and tracking microtubule lattice damage during active transport, complementing existing approaches to studying microtubule integrity [49].

**Figure 3:**
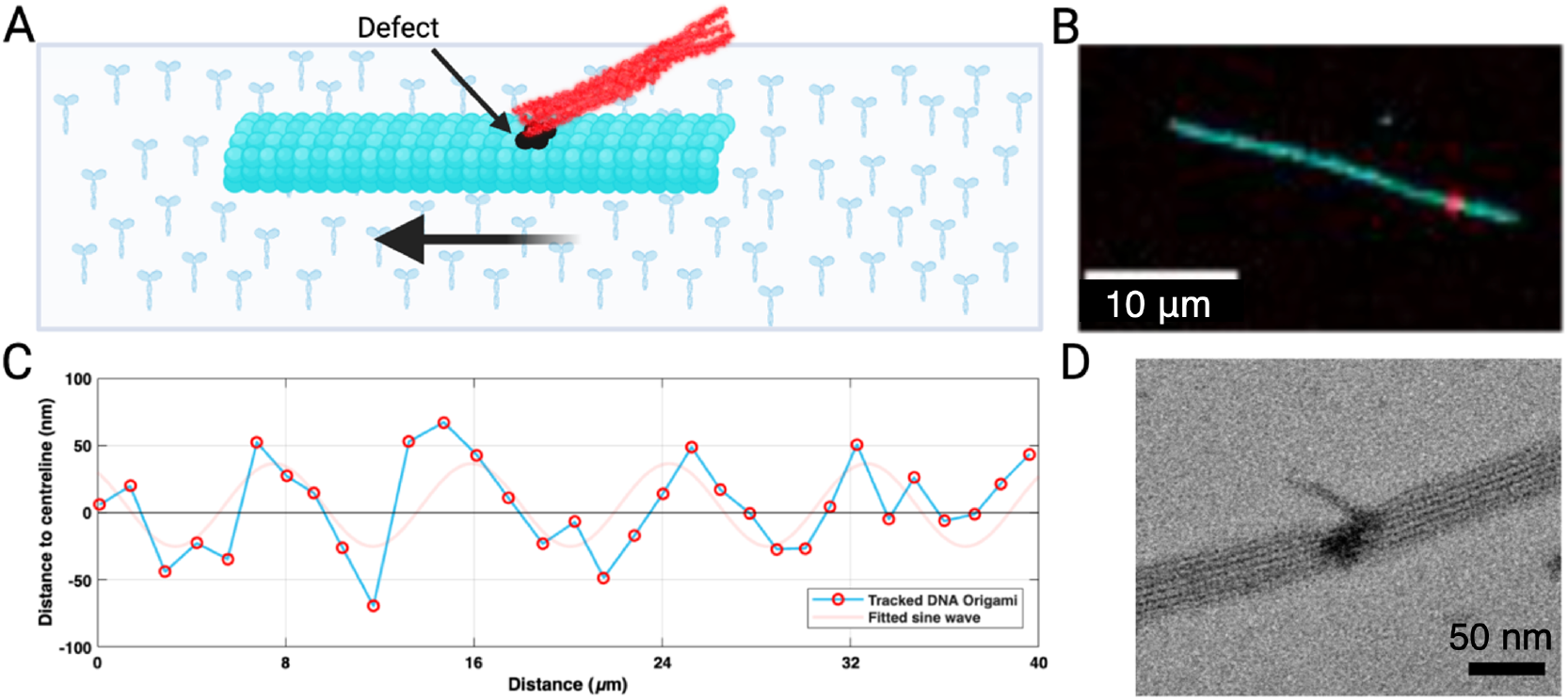
DNA origami caps identify microtubule lattice defects. **A)** Schematic of six-helix bundle labelled with Atto647N (red) attached to a defect site on a TAMRA labelled microtubule (cyan) gliding on surface-immobilized kinesin motors. **B)** Fluorescence microscopy image of TAMRA microtubule (cyan) with Atto647N DNA Origami (red) adhered at a microtubule defect site. **C)** Graph showing DNA Origami at a defect site rotating around a microtubule with a fitted Sine wave in light red of wavelength (pitch) 8.4 μm and Amplitude 30 nm. **D)** TEM image showing six-helix bundle bound to defect site on microtubule (scale bar 50 nm).

To demonstrate functional utility, we investigated whether DNA origami caps could mediate cargo capture during microtubule transport. The six-helix bundle was used as a multivalent platform, with its 22 overhang sequences serving as binding sites for complementary DNA strands. Gliding assays were performed on surfaces containing non-specifically adsorbed Atto647N-labeled oligonucleotides. As capped microtubules moved across the surface, fluorescent signals at their tips increased over time (**Fig. 4A-C**), indicating progressive hybridization and accumulation of oligonucleotides. Maximum intensity projections revealed trajectories that increased in brightness along their path, consistent with continuous cargo pickup. Quantitative analysis confirmed a stepwise increase in fluorescence intensity, approaching the theoretical maximum number of binding sites on the origami. To verify that cargo was acquired from the surface rather than from solution, total internal reflection fluorescence (TIRF) microscopy was employed. Individual surface-bound fluorophores were observed to disappear upon contact with passing capped microtubules (**Fig. 4D; Supplementary Movie 3**), directly confirming pickup from the substrate. This demonstrates that DNA origami caps enable site-specific cargo loading at microtubule ends, distinct from previous approaches based on uniform tubulin functionalization. The system provides a platform for programmable interactions, including cargo collection, transfer, and potentially controlled release through established DNA strand-displacement mechanisms.

**Figure 4:**
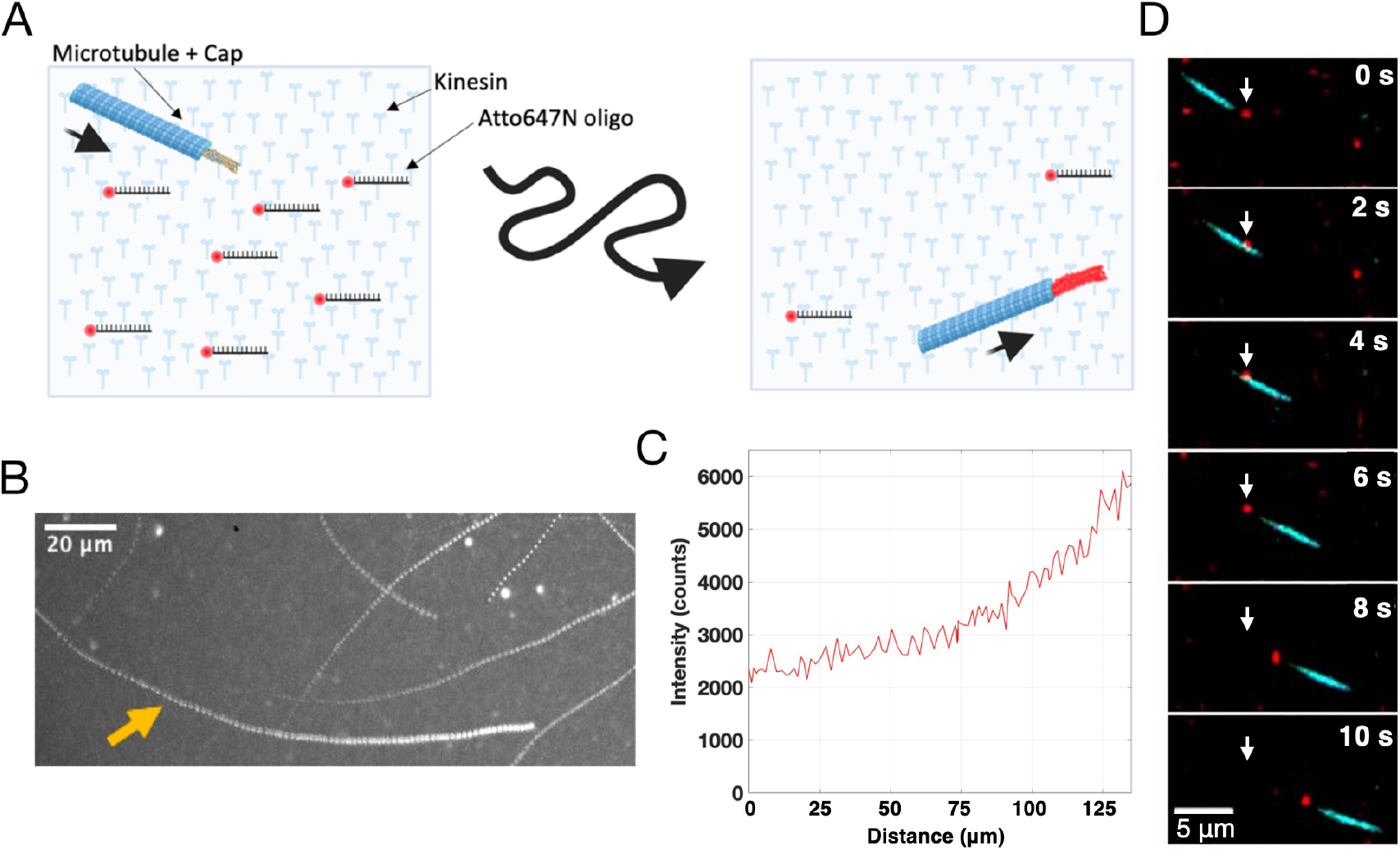
Programmable cargo pickup at microtubule tips during transport. **A)** Illustration showing a microtubule (cyan) with an unlabelled DNA Origami cap (grey) with 22 overhangs for fluorescent labelling gliding on surface-immobilized kinesin motors. As the microtubule with the DNA Origami cap moves over the substrate it picks up non-specifically bound Atto647N oligomers through hybridisation. Thus, the cap and therein the microtubule tip becomes fluorescently labelled. **B)** Maximum projection of a 260 second video of microtubules gliding on an Atto647N coated surface. Several tracks appear to grow brighter as their length increases, for the intensity measurement in **C** the track pointed to by the yellow arrow was used. **C)** Intensity measurement of DNA Origami identified in **B**, tracked using FIESTA. As can be seen the intensity increases over time as the DNA Origami picks up fluorophores from the surface. **D)** TIRF microscopy images of a TAMRA labelled microtubule (cyan) with an unlabelled DNA Origami cap attached to the ‘back’ picking up an individual Atto647N labelled oligomer (red, original position marked by white arrow) non-specifically bound to the surface.

## Discussion

We have introduced a strategy for spatially controlled functionalization of stabilized microtubules by combining luminal antibody targeting with steric exclusion imposed by DNA origami nanostructures. This approach converts a uniformly distributed post-translational modification into a spatially restricted binding modality, enabling selective functionalization at microtubule ends and at lattice discontinuities. In doing so, it addresses a long-standing limitation in microtubule-based nanotechnology, namely the inability to localize functionality to defined positions along the filament. Furthermore, the addition of DNA origami caps does not affect the microtubule gliding velocity, which is typically observed when the cargo binds to the exterior of the microtubules [53], indicating that marking the tips of the microtubules does not hinder interactions between the motors and the microtubules.

The minimal DNA origami six-helix bundle used here demonstrates that precise geometric matching to the microtubule lumen is not required to achieve selectivity. Rather, the combination of lumen-targeting affinity and steric hindrance is sufficient to confine binding to accessible regions. This suggests that a wide design space of DNA nanostructures - varying in size, shape, rigidity, and surface functionality - could be explored to tune binding behavior, valency, and interaction specificity. In this sense, DNA origami caps provide a modular interface through which microtubules can be endowed with programmable and interchangeable functionalities.

One immediate implication of this approach lies in the development of more sophisticated molecular transport systems. Previous strategies for cargo loading in gliding assays rely on uniform functionalization of the microtubule lattice, resulting in distributed binding along the filament. This binding is also associated with a decrease in gliding speed due to hindering of kinesin binding sites [5]. In contrast, the present system enables cargo attachment specifically at microtubule tips and no slowdown is observed. This spatial restriction opens the possibility of directional transport architectures in which cargo is picked up, carried, and delivered in a controlled manner. For example, by combining tip-localized binding with sequence-specific DNA interactions, microtubules could be programmed to selectively collect cargo from defined regions and deposit it at designated target zones. Established mechanisms such as toehold-mediated strand displacement could further enable controlled release, allowing microtubules to function as programmable carriers that execute multi-step transport operations.

Beyond simple pickup and delivery, this framework suggests the possibility of route encoding and history tracking within nanodevices. As demonstrated here, tip-bound DNA origami can accumulate DNA “flags” encountered along a trajectory, effectively recording the path taken by a microtubule. Such a mechanism could be exploited in biocomputational systems or microfluidic networks, where the ability to encode spatial information directly onto moving carriers would provide an additional layer of functionality [50].

The programmable nature of DNA origami also enables interactions between microtubules themselves. Caps could be designed with complementary binding domains that mediate specific end-to-end association, allowing microtubules to fuse into extended filaments or defined architectures. More complex geometries, such as branched junctions, asters, or closed networks, could in principle be assembled dynamically through designed DNA interactions. Such capabilities would extend current approaches to active matter assembly by introducing specificity and reconfigurability at the level of individual filaments.

In addition, the multivalent and addressable surface of DNA origami provides opportunities to modulate microtubule behavior through external control. Caps could be engineered to respond to molecular inputs - such as DNA strands, small molecules, or light - thereby enabling dynamic regulation of binding, cargo interaction, or even motility. For instance, transient interactions between caps and surface-bound DNA could be used to introduce controllable friction or pausing, effectively tuning the speed or dwell time of microtubules within specific regions of a device. Similarly, microtubules could be selectively trapped, released, or redirected through programmable interactions with patterned substrates.

The observation that DNA origami caps also label lattice defects highlights an additional application in the study of microtubule structure and integrity. Microtubules are subject to mechanical stresses and enzymatic activity that can generate lattice damage, yet direct visualization of such defects in dynamic systems remains challenging. The ability of the present constructs to bind at defect sites and report their presence during active transport provides a new tool for probing the formation, distribution, and evolution of lattice discontinuities. This could be particularly valuable for investigating processes such as microtubule aging, repair, and the influence of motor activity on filament integrity. As increasing evidence suggests that defect and repair sites play functional roles - for example as rescue points or as regions of altered biochemical activity - tools enabling their direct observation may offer new insights into cytoskeletal dynamics.

More broadly, the concept of sterically gated access to internal binding sites may be applicable beyond microtubules. Other filamentous or tubular biological structures with internal compartments could potentially be functionalized in a similarly controlled manner by coupling affinity targeting with size-exclusion principles. In this context, DNA origami provides a uniquely adaptable platform for implementing such strategies due to its precise and programmable architecture.

Taken together, our results establish DNA origami caps as a versatile interface between cytoskeletal filaments and synthetic nano systems. While the present work demonstrates proof of concept using a simple six-helix bundle, the range of possible designs and functionalities is substantially larger. By enabling spatially resolved, programmable interactions at microtubule ends, this approach expands the toolkit for motor-driven nanotechnology and opens new directions for the construction of adaptive, reconfigurable, and information-processing nanoscale systems.

## Materials and Methods

### Microtubule and Kinesin preparation

GMP-CPP microtubules overnight at 37 °C from 2 μM porcine tubulin supplemented with 1 mM MgCl_2_ and 1 mM guanosine-5’-[(α,β)-methyleno]triphosphate (GMP-CPP) in BRB80 buffer, pH 6.9 (80 mM 1,4-piperazine-diethanesulfonic acid (PIPES), 1 mM ethylene glycol-bis(β-aminoethylether)-N,N,N′,N′-tetraacetic acid (EGTA), 1 mM MgCl_2_). The microtubules were stabilized and diluted 20-fold in BRB80 containing 10 μM taxol. Full length *Drosophila melanogaster* kinesin-1 motor proteins were expressed in insect cells and purified as previously described [51].

### Flow cell assembly

Flow cells were assembled by placing three parallel strips of parafilm (25×2 mm^2^) spaced apart by ∼2 mm on a 22×22 mm^2^ coverslip (Corning) and a smaller 18×18 mm^2^ coverslip (Corning) on top. Both coverslips were treated with the Easy Clean procedure (sonicated in Mucasol 1:20 for 15 min, washed, and sonicated with Ethanol for 10 min, washed and dried with N_2_). This sandwiched setup was assembled on a thin tissue (Kimtech) and placed on a heating block at 80 °C for ∼15 sec to melt the parafilm strips, thereby creating two imaging channels. The coverslip on top was gently pressed at the sites of the parafilm strips to release any bubbles and ensure leakproof channels.

### Gliding assays in flow cells

A 0.5 mg/ml solution of casein in Brinkley Reassembly Buffer 80 mM (BRB80; adjusted to pH = 6.9 with KOH) composed of 80 mM 1,4-piperazinediethanesulfonic acid (PIPES), 1 mM EGTA, and 1 mM MgCl_2_ was flowed into the cell and incubated for 2 min. Kinesin-1 solution (4 μg/mL kinesin-1, 0.2 mg/mL casein, 10 mM dithiothreitol and 10 μM ATP) was then added and incubated for 10 min. Finally, a motility solution (20 mM D-glucose, 55 μg/mL glucose oxidase, 11 μg/mL catalase, 10 mM dithiothreitol, 10 μM taxol, 1 mM ATP) containing stabilized microtubules (2 μM polymerized tubulin) was added. Excess unbound microtubules and DNA were removed by flushing motility solution without microtubules.

### Sodium dodecyl sulphate – Poly acrylamide gel electrophoresis (SDS-PAGE)

A non-reducing SDS-PAGE was used to analyse the 6-11B-1 antibodies, their fragments and the Fab Complex formation. Samples each containing 0.5 μg of the antibody or its evolutions were mixed with 3 μl of 4x SDS loading buffer and diluted in BRB80 buffer to a total volume of 12 μl. The samples were boiled at 95 °C for 5 min to denature the antibodies and the fragments. 10 μl of the samples were loaded into 4-12% gradient bis-tris gel (Thermo Fisher). 4 μl of an unstained protein marker (PAGE ruler unstained, Thermo Fisher Scientific) mixed with the 4x SDS loading buffer and BRB80 was added to compare the molecular weight. The gel electrophoresis was performed in 1x 2-[N-morpholino] ethanesulfonic acid (MES) running buffer (Thermo Fisher Scientific) at 180 V for 40-45 min. The gels were stained with silver staining to visualize the proteins.

### DNA Origami folding

In order to fold the origami 25 µl of 100nM scaffold (7249 bases; Tilibit) and 6.25 µl of a 4 µM pool of the staple strands were mixed in TE buffer with 6mM MgCl_2_ at a total volume of 50 µl and incubated at decreasing temperature from 95°C (held 1 minute), 80°C (held 10 minutes) ramped from 65°C–45°C (-1.11°C/h) (18 h), 45°C -25°C (-10 °C/h) (2h) and stored at 4°C. The six-helix bundle was then diluted to 500 µl in TE+6mM MgCl_2_ buffer and centrifugated though a 100 kDa MWCO ultracentrifugal filters (MERCK Millipore) at 4 °C and 2000 rcf until 50 µl volume size was reached. The filtrate was discarded, while the remaining mixture containing the six-helix bundle was topped up to 500 μl and spun down for 5 min at 14,000 rcf. This process was repeated three additional times. In the final step, ∼50 μl of six-helix bundle was recovered in a fresh collection vial by the spinning the filters in an inverted position for 1 min at 1000 rcf. Post-purification, origami were quantified using a NanoDrop spectrophotometer (Thermo Scientific) and agarose gel electrophoresis. Azide-modified oligo sequence: /5AzideN/GACGGCTCATAAGGCTCTAATC and Atto647N-modified oligo: TATGAGAAGTTAGGAATGTTA/ 3ATTO647N (645/663nm). The design of the DNA origami is provided in **Fig. S4**.

### Antibody Digestion

The anti-acetyl-α tubulin clone 6-11B-1 monoclonal antibody was treated with the protease papain to digest the full IgG constructs into their Fab fragments. The antibody was supplied in PBS buffer so no initial buffer exchange was required. The pre-activation of papain was performed by treatment of 0.1 mg/ml of crystallized papain solution with 10 mM cysteine and 10 mM EDTA in PBS buffer (pH 7.2) for 10 min at room temperature. 10 μl of the pre-activated papain solution was added to 10 μl of 1 mg/ml IgGs. The digestion reaction was carried out for 1 hour at 39 °C. The reaction was terminated by mixing in 2 μl of 3.3 M iodoacetamide solution for 10 min. For large-scale assays, the reaction was carried out in multiple vials, each containing 10 μl of 1 mg/ml IgG in order to maintain consistency. The Fab fragments were concentrated and purified by diluted to 500 μl in PBS buffer, placed in Amicon 30 kDa MWCO ultracentrifugal filters (MERCK Millipore) and centrifugated for 5 min at 14,000 rcf. The filtrate was discarded, while the remaining mixture containing the Fab fragments was topped up to 500 μl and spun down for 5 min at 14,000 rcf. This process was repeated three additional times. Each cycle results in ∼25 μl of retained volume within the filter resulting in a concentration factor of 20. In the final step, ∼25 μl of Fab fragments were recovered in a fresh collection vial by the spinning the filters in an inverted position for 1 min at 1000 rcf [46].

### Fab Complex Conjugation

25 µl of Fab at an estimated theoretical concentration of 6 µM was added to 1 µl of 6.5 mM DBCO-PEG_4_ -NHSester in DMSO at room temperature for 30 minutes. The excess crosslinker was removed with Zeba Biotin & Dye removal spin column increasing the volume to 50 µl with PBS and spinning through the column at 1000 rcf for two minutes. 20 µl of Fab+crosslinker at an estimated concentration of 3 µM was added to 2 µl of 100 µM azide functionalized oligomer and incubated overnight at 4 °C. The Fab Complex was then purified in Amicon 30 kDA MWCO ultracentrifugal filter with PBS in the same manner as the Fabs were purified above and ∼25 µl recovered.

### DNA Origami-Fab Complex adhesion and Microtubule Capping

Purified Fab Complex was diluted to an estimated theoretical concentration of 150 nM and 3 µl was added to 3 µl of 16.8 nM six-helix bundle and 3 µl of 500 nM Atto647N labelled oligomer or Nuclease free water and incubated at 37 °C for 2.5 hours. 2 µl of Fab Complex-six-helix bundle was then added to 48 µl microtubules in BRB80T and left overnight at room temperature.

### Imaging

Image acquisition was performed using an inverted fluorescence microscope Axio Observer Z1 and a 40× oil α Plan-Apochromat NA = 1.3 objective (Zeiss, Germany) or a 63× oil immersion 1.46NA objective (Zeiss) for TIRF imaging. The data was recorded with an electron multiplying charge-coupled device (EMCCD) camera (iXon Ultra EMCCD, DU-897U, Andor) having a pixel size of 16 μm. Images were acquired with an exposure time of 100 ms using MetaMorph (Molecular Devices, USA).

### Sample preparation and imaging for 2D transmission electron microscopy (TEM)

Carbon-coated 400 mesh copper TEM grids (Merck) were treated with (air) plasma etching for 12 seconds to make the grid surface hydrophilic. Grids were exposed at room temperature to 3-7 μl of sample for 10 min. The sample solution was blotted away, and a droplet of 1% uranyl acetate (Science services) was added to each grid for 30 seconds. Grids were allowed to dry completely for 20 min. The samples were stained for 15 seconds with 1% uranyl acetate (Science services) and imaged at 80 kV on a Jeol JEM 1400 Plus Transmission Electron Microscope. The acquired TEM images were analysed with Fiji.

## Data Analysis

To analyse videos of gliding microtubules on fibre we used in house tracking software FIESTA, built in MATLAB. For determining the speed of microtubules without caps in flow cells we used the AutoTipTrack program [52].

## Acknowledgements

We thank Corina Bräuer for technical support, Krishna Gupta for participating in the origami design, Syuan-Ku Hsiao for help with early conjugation experiments, and Seham Helmi, Qian Zhang, as well as members of the Diez laboratory for scientific discussions. We acknowledge support from (i) the European Union Marie Skłodowska-Curie Actions Innovation Training Network (ArtMoMa) and the Dresden International Graduate School for Interdisciplinary Life Sciences (DIGS-ILS) for H.C.M., (ii) the Deutsche Forschungsgemeinschaft (DFG) within GRK 2767: Supracolloidal Structures: From Materials to Optical and Electronic Devices (Project No. 451785257) for B.P.R., and (iii) the Federal Ministry of Research, Technology, and Space (BMFTR) in the program NanoMatFutur (grant no. 13XP5098) for E.K..

## Author Contributions

H.C.M., B.P.R. and S.D. conceived and designed the study. H.C.M. and B.P.R. performed the research and analyzed the respective data. A.A., J.B., A.T., E.K. contributed reagents and technologies. H.C.M. and S.D. wrote the manuscript with contributions from the other authors. All authors discussed the results and approved the final version of this manuscript.

