## Supplementary Figures for "Programmable DNA Origami Caps for Site-Selective Functionalization of Microtubule Tips and Lattice Defects"

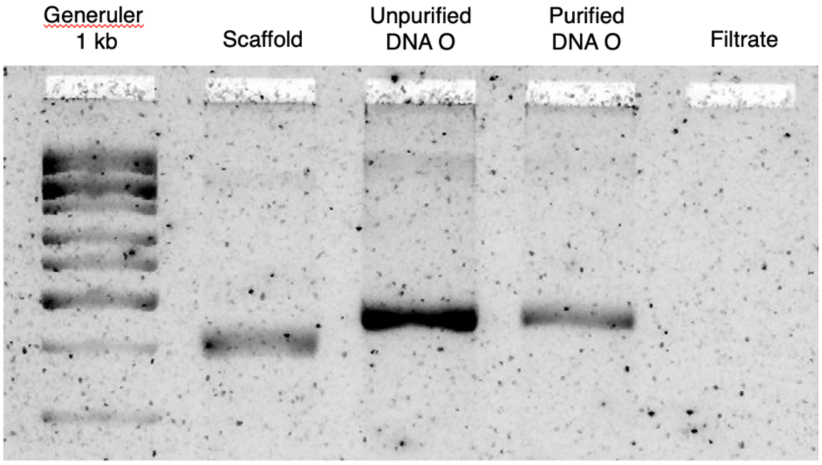


***Figure S1: DNA Origami folding and Agarose Gel Electrophoresis.*** *1.5% Agarose gels were prepared with high melt agarose (Invitrogen) and 1x TBE 1 µl of scaffold at 0.1X, 1 µl of unpurified 6-helix bundle, 1 µl of purified 6-helix bundle and 8 µl of filtrate were mixed 2 µl 10X Bluejuice loading buffer (BLD) were loaded onto the gel. 4µL Genruler 1 kb (500 ng) + 1 µL BLD was used to compare molecular weight. The separation took place in (1x TBE + 11mM $MgCl_2$) running buffer for two hours at 60 V. The gel was stained with DNA SYBR Gold for half an hour and imaged in a Typhoon Gel Dock.*

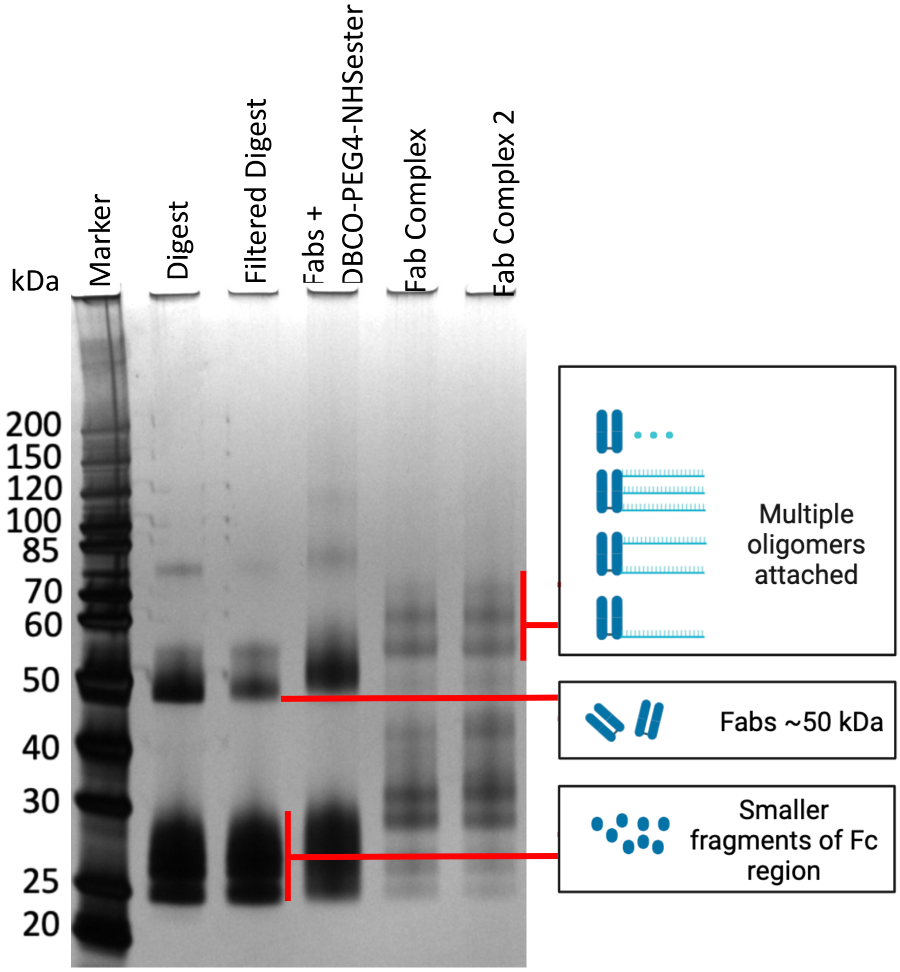


***Figure S2: Additional SDS-PAGE showing multi-oligomer Fab Complex formation.*** *6-11-B1 antibodies were digested with papain and then incubated with the DBCO-PEG$_4$-NHS ester crosslinker. The sample was then halved with one having excess crosslinker removed using 30 kDA (Merck Millipore) column and the other with a Zeba Dye and Biotin desalting column as per the standard protocol. The two were then separately incubated with the azide functionalised oligomer which reliably causes separation in the gel into multiple bands depending on the number of oligomers attached.*

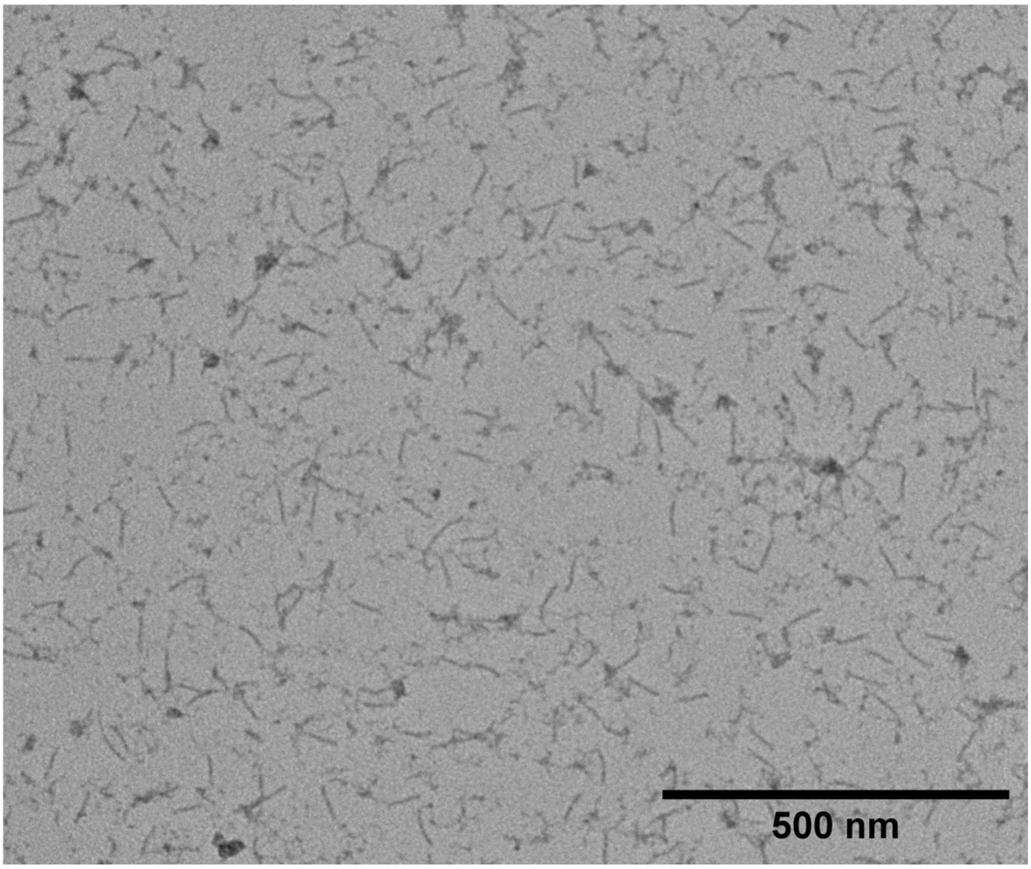


***Figure S3: TEM image of Fab Complex 6-helix bundle caps.*** *See Methods section for sample preparation and imaging.*

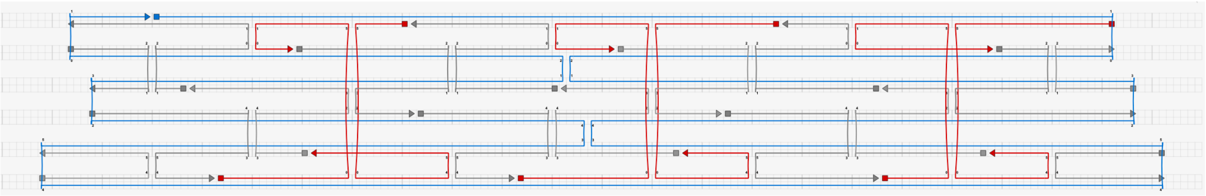

***Figure S4: CaDNAno [1] design scheme of the mini-6HB used in this study.*** *The folded origami structure comprises approximately 900 base pairs, while the remainder of the 7249-nt scaffold remains single-stranded by design.*
